# Results of follow-up experiments to “Odorant cues linked to social immunity induce lateralized antennal stimulation in honey bees (*Apis mellifera* L.)”

**DOI:** 10.1101/232249

**Authors:** Alison McAfee, Troy F. Collins, Leonard J. Foster

**Affiliations:** Department of Biochemistry & Molecular Biology and Michael Smith Laboratories, University of British Columbia, 2125 East Mall, Vancouver, British Columbia, Canada

## Abstract

In 2017, we published the paper “Odorant cues linked to social immunity induce lateralized antennal stimulation in honey bees (*Apis mellifera* L.)” in *Scientific Reports*. Since then, we have performed three follow-up experiments which have either negative or contradictory results. Previously, we used electrophysiology to show that hygienic bees displayed significantly higher sensitivity to β-ocimene when stimulated via their left antennae compared to their right. We repeated this assay using worker honey bees from a single hygienic colony and found, to our surprise, that the right antennae elicited higher sensitivity. We also previously attempted to identify a molecular basis for lateralization by using mass spectrometry-based proteomics to compare left and right antennal proteomes. Of the 1,845 proteins, none were differentially expressed. Here, we repeated this experiment but employed orthogonal peptide fractionation to increase proteome coverage to 3,114 proteins; however, still none were differentially expressed. Finally, we attempted to manipulate gene expression of a key antennal odorant binding protein linked to hygienic behaviour (OBP18) using RNA interference via antenna microinjection. We were not able to achieve long-lasting OBP18 knock-down, but comparing the proteomes of untreated, mock dsRNA-treated and OBP18 dsRNA-treated worker antennae revealed numerous off-target effects of the act of injecting alone. By openly reporting this data, we hope to set an example for information transparency.

## Introduction

Publication bias – most commonly, the tendency for journals to publish only significant and novel results – is a well-known pitfall of publishing scientific data in conventional peer reviewed journals (1, 2). This can be to the detriment of science, since negative results (those showing no statistical significance) are no less true than positive results (which do show statistical significance), and are often still worthy of dissemination. Publication bias also selects against replication experiments, so results are commonly not directly corroborated by additional studies (3). Here, we openly publish three key pieces of negative or contradictory data that directly follow one of our previous publications.

Recently, we published the paper “Odorant cues linked to social immunity induce lateralized antennal stimulation in honey bees (*Apis mellifera* L.)” in *Scientific Reports* (4). Since then, we have followed up on our electroantennography results and found that our new data are inconsistent with what we published previously. We also previously performed a quantitative proteomics experiment to try to elucidate a mechanism to explain antennal lateralization. When we did not identify any differentially expressed proteins, we reasoned that it might be because of poor proteome coverage. Therefore, we repeated our proteomics experiment with greater depth of coverage, but still found no statistically significant results. Finally, we attempted to manipulate antennal expression of OBP16 and OBP18 – proteins linked to social immunity (5, 6) – using RNA interference, but we were unable to achieve long-lasting knockdown. Although these results may not be significant, we think this information highlights the importance of replication studies and, importantly, the microinjection-induced off-target effects may assist researchers who are attempting to use a similar approach.

## Results

Previously, we used electroantennography (EAG) to measure amplitudes of the collective antennal nerve depolarizations of honey bees when presented with an odor (4). Specifically, we compared the left and right antennae of hygienic and non-hygienic bees upon stimulation with β-ocimene – a compound which is strongly emitted from freeze-killed brood. Our previous experiments identified a significant interactive effect between side and hygienic behaviour; however, the study suffered from relatively low replication (N = 10-15 antennae per group), so we sought to confirm the results with a more robust sample.

In a subsequent experiment, we compared the left and right antennal response to β-ocimene using N = 22 bees from a single highly hygienic colony. We expected to observe the same left-biased response as before; however, we observed the opposite pattern (Figure 1). We still observed a significant dose-response (two-way ANOVA; F = 27.5, p = 1.5e-10), but the right antennae responded marginally (but significantly) more strongly than the left (F = 6.3, p = 0.01).

**Figure 1.**
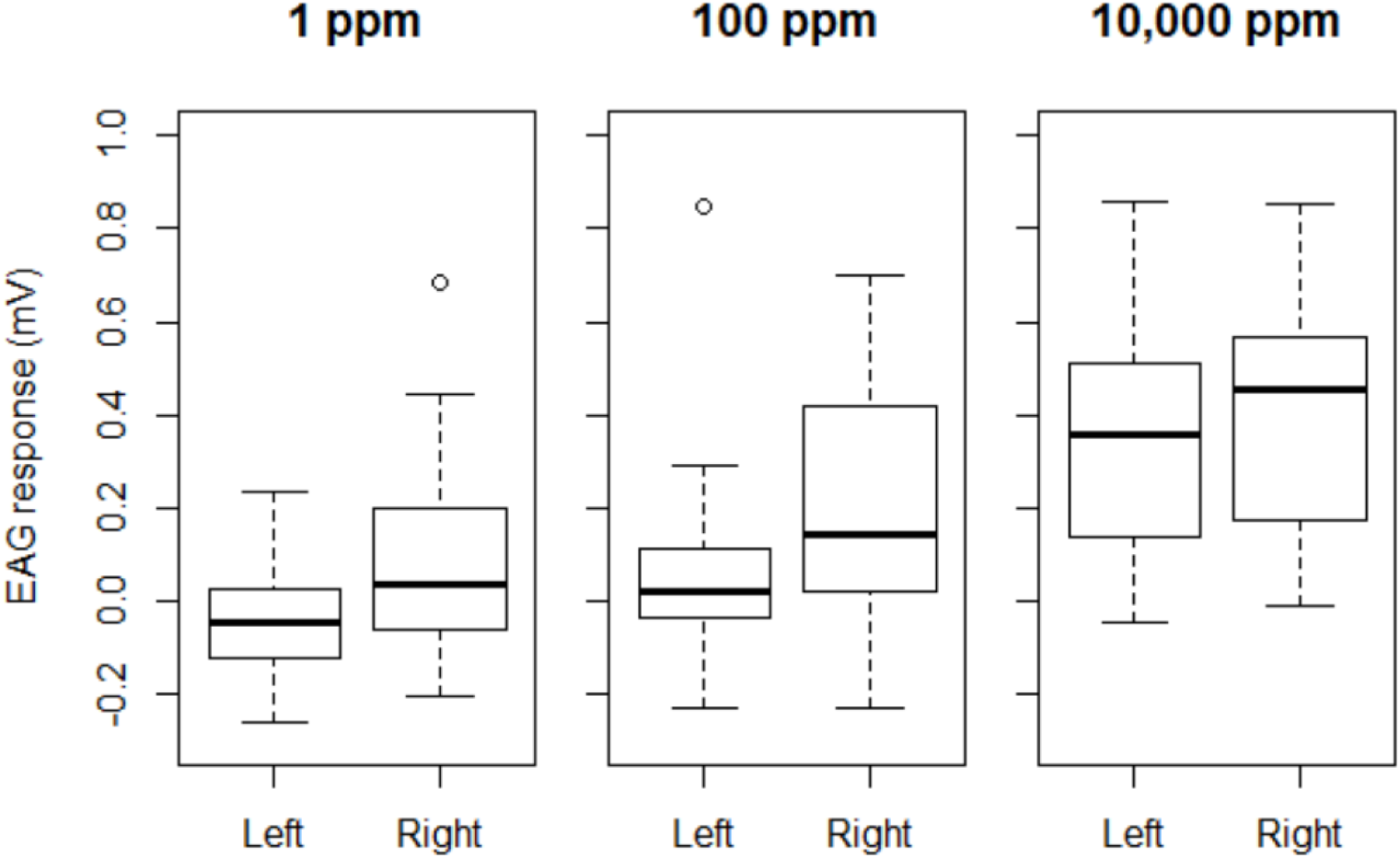
*Electroantennography (EAG) responses to β-ocimene.* Bees’ antennae from a single hygienic colony (hygienic score: 95%) were excised and stimulated with three increasing concentrations of β-ocimene (N = 22). β-ocimene was serially diluted in ethanol. The plotted EAG response is corrected for the background stimulation of ethanol alone. Data was analyzed with a two-factor ANOVA (levels: side and dose). There was a significant effect of dose (F = 27.5, p = 1.5e-10) and side (F = 6.3, p = 0.01). Boxes depict the interquartile range (IQR), whiskers span 1.5*IQR, and bold bars represent the median.

Previously, we attempted to identify the molecular basis for lateralization by comparing protein expression between left and right antennae of hygienic bees using mass spectrometry-based quantitative proteomics (4). We identified 1,845 unique proteins but found no significant differences; therefore, we suggested this may be because of insufficient depth of coverage. To improve proteome coverage in the current study, we fractionated the peptides from left and right antennae from 4 highly hygienic colonies and repeated the comparison (Figure 2). This time, we identified 3,114 unique proteins (a 69% improvement), which is among the highest proteome coverage achieved in honey bees to date (7). However, we still did not identify significant differences. Hierarchical clustering shows that the data groups by colony, rather than by left or right.

**Figure 2.**
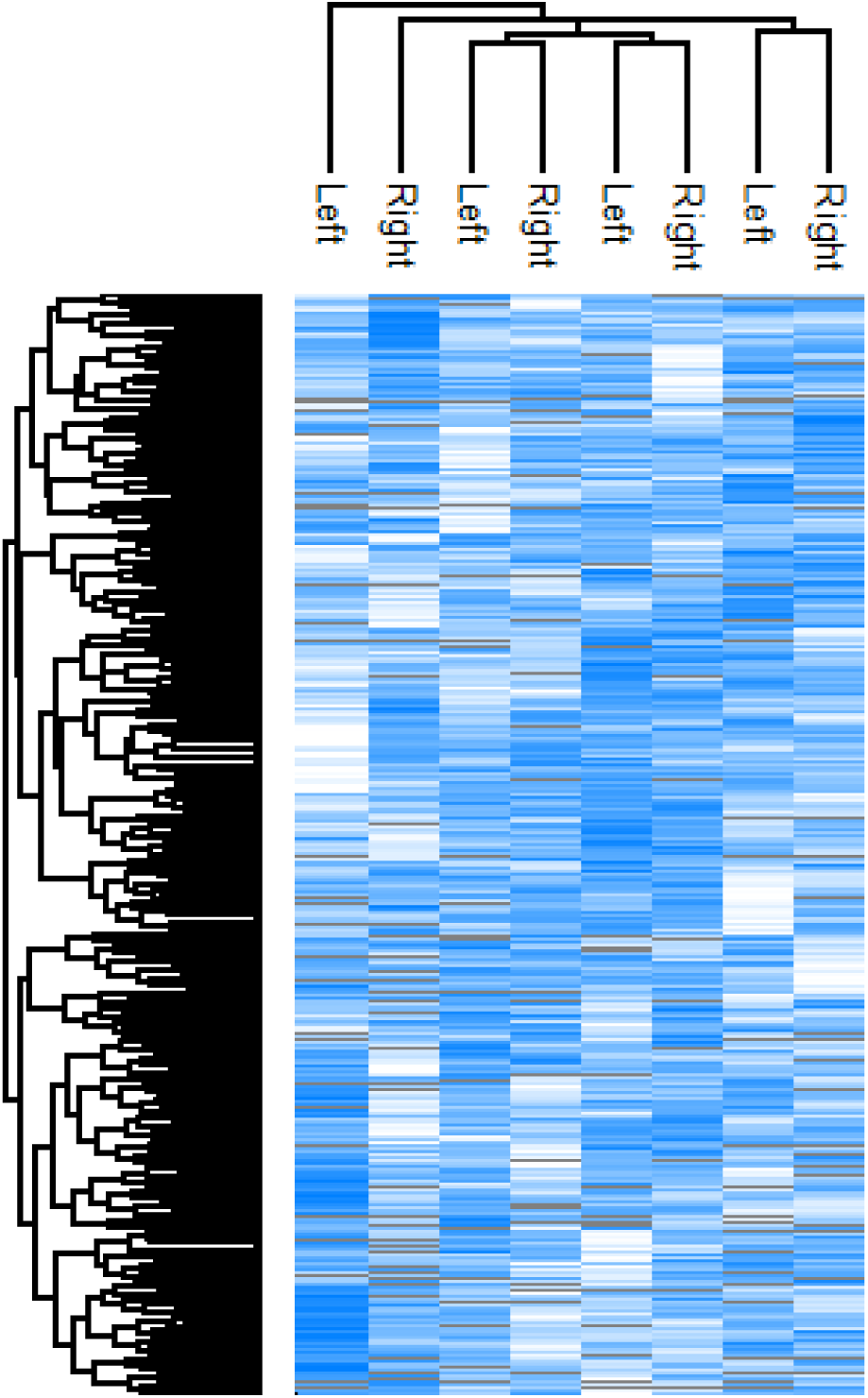
*Comparing the proteomes of left and right antennae from hygienic honey bees.* Digested peptides were fractionated by basic reverse phase chromatography (5 fractions each) and analyzed on a Bruker Impact II Q-TOF mass spectrometer. Label-free quantitation was used to compare protein expression between samples. 3,114 proteins were identified, but no proteins were significantly different. Z-score scale: white = −2.5, blue = +2.5, and grey = not quantified. Hierarchical clustering was performed in Perseus using average Euclidian distance (300 clusters, maximum 10 iterations). Raw files and protein identifications can be found at ftp://massive.ucsd.edu/MSV000081790.

Previous studies have found that OBP16 and OBP18 expression is significantly positively correlated with hygienic behaviour. Odorant binding proteins such as these are thought to be important for binding airborne odorants and transporting them to the olfactory receptors, stimulating nerve depolarization (8, 9). Since we already found some candidate hygienic behaviour-inducing odors, we wished to experimentally manipulate the OBPs and see if this changed the bees’ responses to these odors. We therefore attempted to knock down OBP16 and OBP18 using RNA interference by directly microinjecting the dsRNA into the antennae of pupae. We used targeted mass spectrometry (multiple reaction monitoring, or MRM) to quantify unique peptides from these proteins in untreated, mock treated, and OBP-specific dsRNA-treated pupa antennae. However, we were unable to confirm a knock-down response, since even GFP (mock) dsRNA appeared to induce equivalent knock-down as the dsRNA that was meant to target OBP18 (Figure 3). Furthermore, while OBP18 dsRNA appears to target OBP18 better than OBP21 (which has a similar sequence), this effect is weak at best and is lost when the dsRNA undergoes the supplier’s (AgroRNA’s) proprietary purification method. OBP16 was not detectable in any of the samples – even the untreated control – suggesting that it is not expressed in newly emerged adults at all.

**Figure 3.**
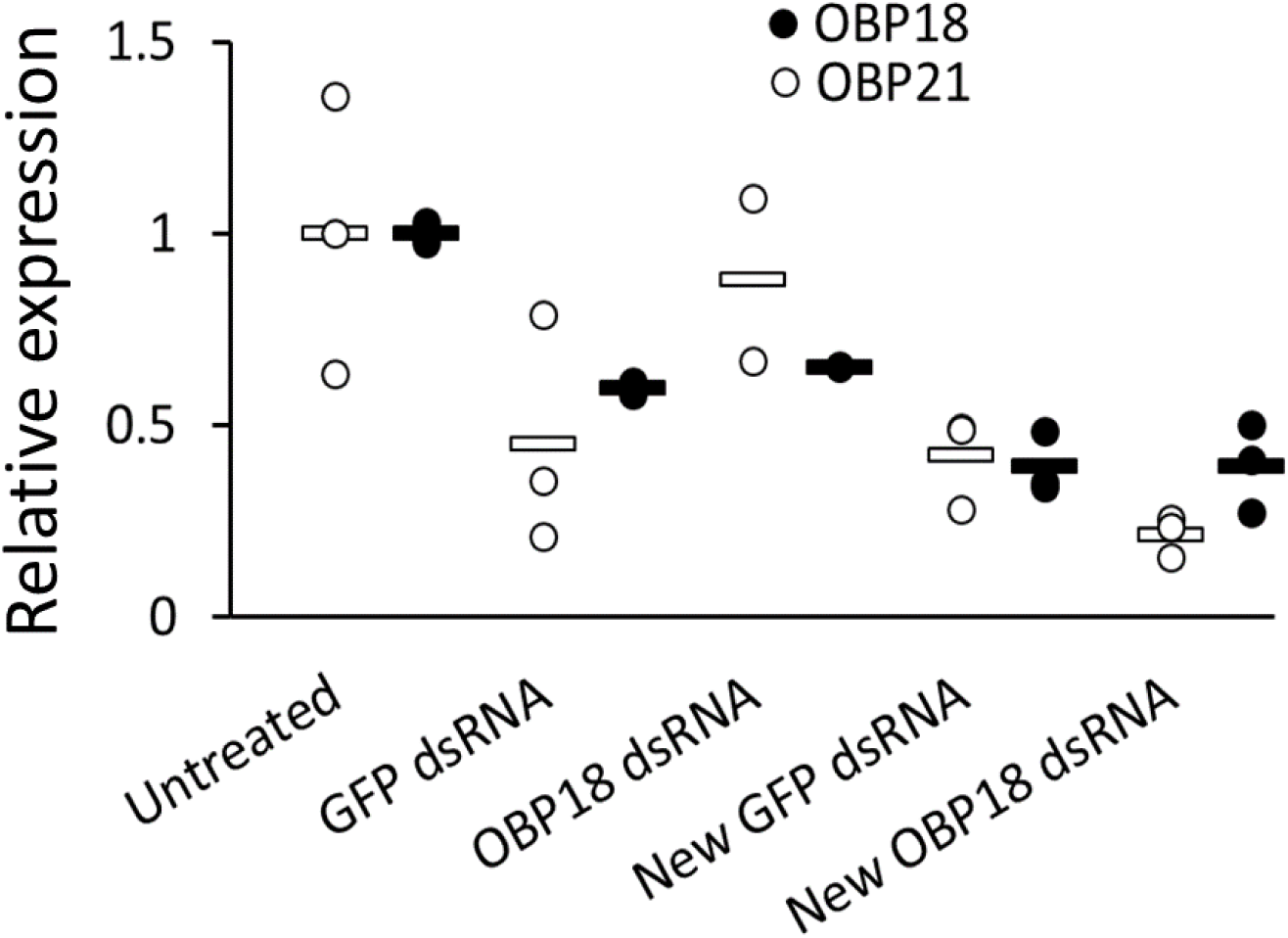
*RNA interference against OBP18 in the antennae.* We microinjected 3 ng of long dsRNA (GFP: 252 bp, OBP16: 448 bp, OBP18: 453 bp) into the flagella of pink-eyed pupae and allowed them to reach adulthood before harvesting their antennae. We then extracted the antennal proteins and quantified OBP18 and OBP21 peptides using multiple reaction monitoring mass spectrometry (Agilent 6460 QQQ). We were unable to detect endogenous OBP16, even in untreated samples. We used antennae from 10 bees per sample, and each treatment group has 3 replicates (except OBP18 dsRNA, which is in duplicate). Untreated bees received no dsRNA, GFP and OBP18 dsRNA bees received the GFP or OBP18 sequence (synthesized by AgroRNA but purified in-house), and new GFP and OBP18 dsRNA bees received the same GFP or OBP18 dsRNA molecule, but which underwent a proprietary purification method by AgroRNA. dsRNA sequences are available in Supplementary information.

Finally, we performed a shot-gun proteomics analysis on the same protein preparation used for the above MRM experiment, and found that simply the act of microinjecting had a profound effect on protein expression (Figure 4). Some 2,852 proteins were identified at 1% FDR (2,586 quantified by LFQ), 918 of which were significantly different between treatment groups (one-way ANOVA, levels: untreated, OBP18 dsRNA, GFP dsRNA; Benjamini-Hochberg corrected at 5% FDR).

**Figure 4.**
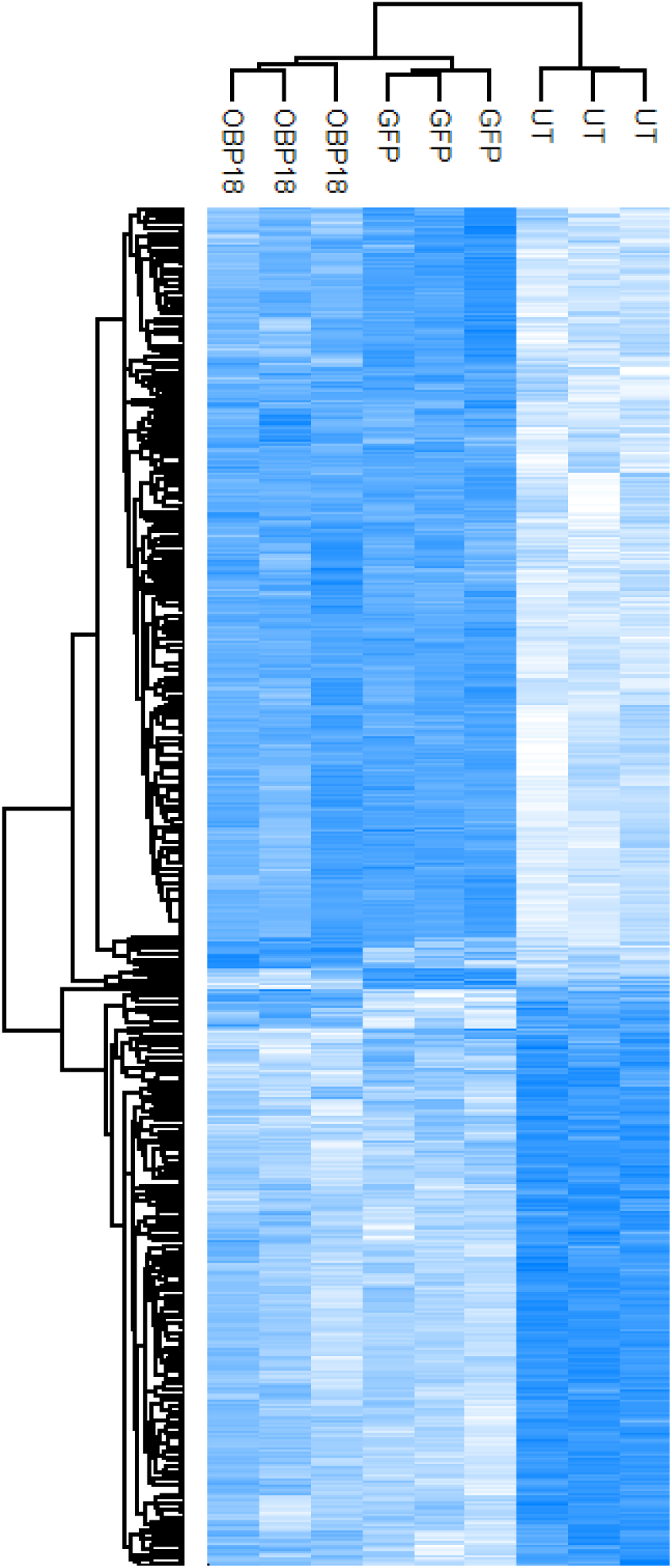
*Heatmap of shot-gun proteomics analysis comparing microinjected antennae.* We identified 2,852 proteins (25,640 unique peptides, 1% peptide and protein FDR) using a Bruker Impact II QTOF mass spectrometer coupled to an EASY-nLC 1000. This heatmap shows only the 918 proteins that were differentially expressed (label-free quantitation, one-way ANOVA, Benjamini-Hochberg corrected 5% FDR). OBP18 was not one of them. Z-score scale: white = −2.5, blue = +2.5, and grey = not quantified. Hierarchical clustering was performed in Perseus using average Euclidian distance (300 clusters, maximum 10 iterations). UT = uninjected. OBP18 = antennae were injected with OBP18 dsRNA. GFP = antennae were injected with GFP dsRNA. Raw files and protein identifications can be found at ftp://massive.ucsd.edu/MSV000081790.

## Discussion

Here, we present the results from three follow-up experiments to our previous publication, “Odorants linked to social immunity induce lateralized antennal stimulation in the honey bee (*Apis mellifera* L.)” (4). First, we show that the lateralized response to β-ocimene (biased to the left antenna) is not consistent, since we could not reproduce our earlier findings when we repeated the experiment the following year. Instead, we found that the response is right-biased (Figure 1). Regardless of directionality, we also sought to find the molecular mechanism of lateralization by improving the depth of our previous quantitative proteomics experiment; however, we still did not detect significant differences between left and right antennae of hygienic honey bees (Figure 2). Finally, we wished to investigate the relationship between the disease odors we identified in our previous publication and OBP18 – a protein biomarker for hygienic behaviour – using RNA interference. Unfortunately, we were unable to achieve consistent knock-down, even by directly microinjecting into the honey bees’ antennae (Figure 3), and found significant off-target effects on protein expression just from injecting mock dsRNA (Figure 4).

It’s unclear why the lateralized response to β-ocimene is not consistent. It’s possible that in our previous experiment, the low sample size (between 10 and 15 antennae, depending on the treatment group) led us to draw incorrect conclusions. Handedness has recently been shown to be important in honey bee (10) and bumble bee (11) behaviours. If handedness is also linked to antenna lateralization, this would make having a large sample size for EAG even more critical.

Left and right antennae have different morphology and responsiveness to odors (12) – differences which likely have underpinnings in gene expression. Despite improving our proteome coverage by 69%, we still could not identify any differential protein expression between left and right antennae. As we stated previously, it is still possible that the lateralization shift is caused by very low abundance proteins (indeed, 3,114 proteins corresponds to only about 20% of annotated honey bee genes (13)). However, it is also possible that differentially spliced proteins could contribute to lateralization, which we could not distinguish between. It is also possible that differences in protein expression are stronger during antennal development (*i.e.* within pupae) and the differences are not as apparent in adults, which is the only stage we analyzed.

RNA interference is conceptually simple and works well as a molecular biological tool in many organisms. Several studies in insects have reported that RNAi induced by long dsRNA can elicit longterm, systemic knock-down responses (14-16). However, we found that even direct injection of long dsRNA into the antennae of pupae could not elicit an RNAi response (measured after 1 week) against OBP18. We have tried other methods of dsRNA delivery, including larval feeding, adult feeding, and aerosolized lipid nanoparticles; however, all of them yielded similarly unconvincing results. It could be that systemic RNAi is not readily achievable in honey bees – indeed, one of the only examples of systemic RNAi in honey bees is from an experiment where the dsRNA corresponds to a honey bee virus, rather than the bees’ own genes (17). We are therefore pursuing transgenics as a method of manipulating gene expression in honey bees (18), with the ultimate goal of determining a relationship between OBP expression and death or disease odor detection.

We also attempted to perform RNAi against OBP16 (another OBP linked to social immunity). Using targeted mass spectrometry, we monitored two peptides corresponding to OBP16; however, we were unable to detect them in the antennae of newly emerged adults. Foret *et al.* (8) determined that OBP16 was not expressed in old pupae, but it is expressed in foraging adults. Previous studies in our own lab suggest that it is expressed in nursing adults as well. However, since we did not detect OBP16 even in our untreated samples, this suggests that newly emerged adults are too young to have activated OBP16 expression.

## Conclusion

Although these experiments yielded either non-significant or contradictory results, they are important for two reasons. First, we need to encourage reporting of replication studies, regardless of the results, to improve data reliability. Second, even negative results can be important experimental outcomes that can guide the direction of future experiments. By reporting the outcomes of three experiments that directly followed our previous peer-reviewed publication, we hope to improve transparency and expose data that would normally go unpublished.

## Methods

### Honey bee samples

All honey bees were obtained between summer 2016 and the spring of 2017. Colonies were kept at different locations within the Greater Vancouver Area: our rooftop apiary at the University of British Columbia, at the UBC farm, and in Abbotsford. Hygienic testing was performed as previously described (19). Honey bees for electroantennography and antennal shotgun proteomics analysis were obtained from open brood frames within a single hygienic (FKB score = 95%) colony. Honey bee pupae for microinjections were sampled from capped brood frames using forceps.

### Electroantennography

Electroantennography experiments were performed exactly as previously described (4). All statistical analyses were performed in R unless otherwise specified.

### Deep antennal proteomics

We dissected approximately 30 pairs of worker bee antennae from each of 4 highly hygienic colonies. Proteins were extracted and processed for mass spectrometry exactly as previously described (4), except after digesting 30 μg of protein, the peptides were fractionated using basic reverse phase chromatography (20). We pooled every 6^th^ fraction, dried them down (Eppendorf Speed Vac), and acidified them in 0.5% formic acid prior to loading 20% of the sample (approximately 1 μg) on a Bruker Impact II QTOF mass spectrometer (coupled to a Thermo EASY-nLC 1000 chromatography system) for shotgun proteomics analysis. The liquid chromatography and mass spectrometry parameters and specifications were set as previously described (4). Data was processed and analyzed by label-free quantification (MaxQuant v1.5.3.30) and statistical analysis was performed in Perseus (v1.5.5.3) as previously described (21). All proteomics data and protein identifications can be found at ftp://massive.ucsd.edu/MSV000081790.

### RNA interference

We used long (> 200 bp) dsRNA corresponding to GFP (252 bp), OBP16 (448 bp), and OBP18 (453 bp) for the RNAi experiments. All dsRNA was synthesized by AgroRNA (Seoul, South Korea). Sequences are available in the Supplementary Information. OBP16 and OBP18 sequences include only unique gene regions (>50 bp) to avoid off-target effects with similar OBPs. The short unique regions were concatenated to yield a longer dsRNA, since evidence suggests they are more effective than short dsRNA molecules (15). The dsRNA was either purified by isopropanol precipitation upon receipt from the manufacturer, or the manufacturer performed a proprietary purification (indicated by “New” dsRNA in Fig. 3). Integrity of the dsRNA was always confirmed by gel electrophoresis prior to commencing RNAi experiments. We also confirmed the activity of the GFP dsRNA by transfecting *Drosophila melanogaster* S2 cells with a GFP expression plasmid (X-tremeGENE HP, manufacturer’s protocol), followed by either mock or GFP dsRNA treatment by media soaking. GFP expression was assessed by a Western blot (Figure S1).

To perform RNAi *in vivo* in honey bees, we microinjected 3 ng of purified dsRNA (suspended in nuclease-free water) into the flagella of pink-eyed worker pupae. We kept the pupae in a humid incubator (33°C) until they reached adulthood (1 week), at which time we euthanized them with CO_2_ and dissected their antennae (10 pairs per replicate) for shotgun proteomics. Samples were processed as described above, except no basic reverse phase fractionation was performed. Raw data and protein identifications can be found at ftp://massive.ucsd.edu/MSV000081790.

### Multiple reaction monitoring

Protein was extracted from the antennae, quantified, reduced, alkylated, and digested with trypsin as previously described (21), except heavy stable isotope labelled peptide standards for OBP16 and OBP18 were added prior to reduction. No standards were used for OBP21 peptides. Peptides were desalted using a high-capacity C18 STAGE tip prior to mass spectrometry analysis. Multiple reaction monitoring (MRM) was performed by injecting 8 μg of digested protein as previously described (19). See Table 1 for a list of peptides and transitions monitored.

**Table 1.** Peptides analyzed by MRM.

| Peptide sequence<sup>1</sup> | Protein | fmol<br>peptide/µg<br>protein | Transitions<sup>2</sup> |
| --- | --- | --- | --- |
| EIAEIYLDENEV NK | OBP21 |  | 839.9(+2) → **960.5(+1)**, 1123.5(+1), 1436.7(+1) |
| NGIIDVENEK | OBP21 |  | 565.8(+2) → 618.3(+1), **733.3(+1)**, 846.4(+1) |
| TGIQTLQPICVGETGTSQK | OBP16 |  | 673.3(+3) → 638.8(+1), 742.4(+1), **807.4(+1)** |
| TGIQTLQPICVGETGTSQK<sup>3</sup> | OBP16 | 200 | 676.0(+3) → 642.8(+1), 742.4(+1), **815.4(+1)** |
| TITDILNS | OBP16 |  | 438.7(+2) → 771.4(+1), **657.4(+1)**, 544.3(+1) |
| TITDILNS<sup>4</sup> | OBP16 | 200 | 442.2(+2) → 778.4(+1), **664.4(+1)**, 551.3(+1) |
| EIAEIFLDENG VNK | OBP18 |  | 795.9(+2) → 775.4(+1), **1035.5(+1)**, 1148.6(+1) |
| EIAEIFLDENG VNK | OBP18 | 100 | 799.9(+2) → 783.4(+1), **1043.5(+1)**, 1156.6(+1) |
| IETSIDQQK | OBP18 |  | 531.3(+2) → 518.3(+1), 718.4(+1), **819.4(+1)** |
| IETSIDQQK | OBP18 | 100 | 535.3(+2) → 526.3(+1), 726.4(+1), **827.4(+1)** |
| DGNIDVEDEK | OBP18 |  | 567.3(+2) → 619.3(+1), **734.3(+1)**, 847.4(+1) |
| DGNIDVEDEK | OBP18 | 100 | 571.3(+2) → 627.3(+1), **742.3(+1)**, 855.4(+1) |

1. Heavy residues within the peptide sequence are in bold 2. Only bold transitions were used for quantification 3. The underlined cysteine was reduced and alkylated during peptide synthesis 4. This is a C-terminal peptide; therefore, an internal residue (I) was labeled

## Supplementary information

### GFP dsRNA sequence

ATGGTGAGCAAGGGCGAGGAGCTGTTCACCGGGGTGGTGCCCATCCTGGTCGAGCTGGACGGCGACGTAAACG
GCCACAAGTTCAGCGTGTCCGGCGAGGGCGAGGGCGATGCCACCTACGGCAAGCTGACCCTGAAGTTCATCTGCA
CCACCGGCAAGCTGCCCGTGCCCTGGCCCACCCTCGTGACCACCCTGACCTACGGCGTGCAGTGCTTCAGCCGCTA
CCCCGACCACATGAAG

### OBP16 dsRNA sequence

GTACAATTTAAACTATAAAAAGATCTTGTACATTCGACGTTTCTTGAGTTTTCCTTAAAAATATTTCAAGAACGTACA
ATtTATCTCTGATGCTGACTTAGCTGTAAAATCTGCTAAATTATTGAAGTGTATTGGAAAATGTACAATTATCTCTGA
TGCTGACTTAGCTGTAAAATCTGCTAAATTATTGAAGTGTATTGGAAAATGTACAATTATCTCTGATGCTGACTTAG
CTGTAAAATCTGCTAAATTATTGAAGTGTATTGGAAAATGTACAATTATCTCTGATGCTGACTTAGCTGTAAAATCT
GCTAAATTATTGAAGTGTATTGGAAAATGTACAATGTAACACAATATTTTTTCTTTATTTTAAAATTGTTTTAATTAT
ACTTTGATTATAATTATATTAATTATACTTTATTATTATACTTTTAACTTTTATTATATGTACAAT

### OBP18 dsRNA sequence

GTACAATGCATATTCGATATTTGTTCAGTTCTCGTTGAACGTTTCAAGAATAGTCGAGTATTTTTATTTATTTGAAAT
CGAtGTACAATGTTGGTGCAATGACACATGAGGAATTAAAAACCGGAATACAGACTTTACAGCCAATTTGCGTAGG
CGAAACTGGCACTAGTCAAAAAATAATAGATGAAGTTTATAATGGCAACGTCAATGTAGAAGACGAAAATGTGTA
CAATGTTGGTGCAATGACACATGAGGAATTAAAAACCGGAATACAGACTTTACAGCCAATTTGCGTAGGCGAAAC
TGGCACTAGTCAAAAAATAATAGATGAAGTTTATAATGGCAACGTCAATGTAGAAGACGAAAATGTGTACAATTA
AAATATATTTGAAAACTTTTATTAATAAATCAATATTATATATTATTATAAATTAATTATTAGTACAAT

---

**Figure S1.**
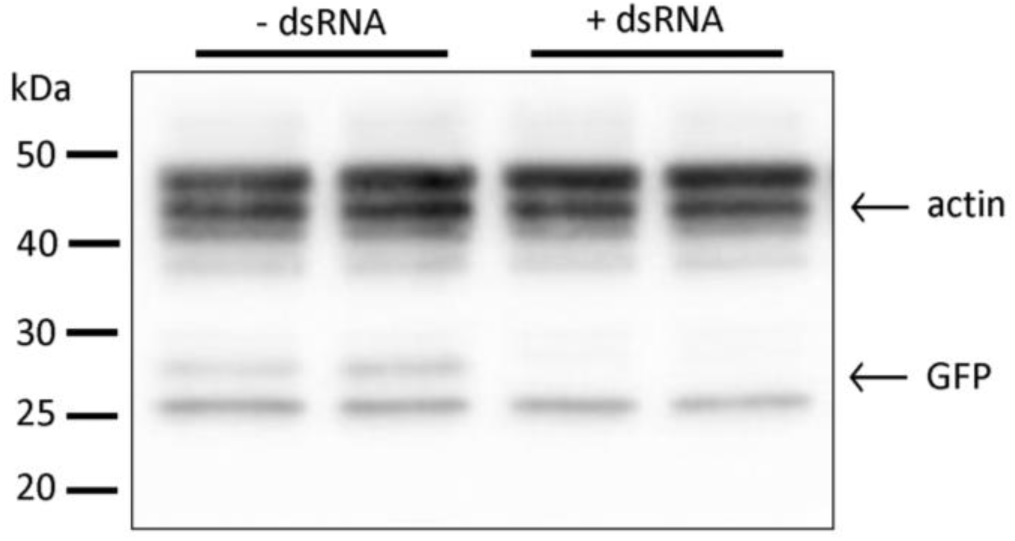
*Western blot confirming dsRNA activity*. *Drosophila* S2 cells were transfected with a GFP expression plasmid followed by either mock (replace media, no dsRNA) or GFP dsRNA treatment via media soaking. We loaded 20 μg of protein in each lane and co-probed for GFP (27 kDa) and actin (42 kDa).

